# Proteomic analysis of mouse kidney tissue associates peroxisomal dysfunction with early diabetic kidney disease

**DOI:** 10.1101/2021.10.21.465240

**Authors:** Aggeliki Tserga, Despoina Pouloudi, Jean Sébastien Saulnier-Blache, Rafael Stroggilos, Irene Theochari, Harikleia Gakiopoulou, Harald Mischak, Jerome Zoidakis, Joost Peter Schanstra, Antonia Vlahou, Manousos Makridakis

## Abstract

**Background:** The absence of efficient inhibitors for DKD progression reflects the gaps in our understanding of DKD molecular pathogenesis. A comprehensive proteomic analysis was performed on glomeruli and kidney cortex of diabetic mice with subsequent validation of findings in human biopsies and - omics datasets aiming to better understand the underlying molecular biology of early DKD development and progression.

**Methods:** LC–MS/MS was employed to analyze the kidney proteome of DKD mouse models: Glomeruli of Ins2Akita mice 2 month and 4 month old, and cortex of db/db mice 6 month old. Following label-free quantification, the abundance of detected proteins were correlated with existing kidney datasets and functionally annotated. Tissue sections from 16 DKD patients were analyzed by IHC.

**Results:** Pathway analysis of differentially expressed proteins in the early and late DKD versus controls predicted dysregulation in DKD hallmarks (such as peroxisomal lipid metabolism, β-oxidation and TCA cycle) supporting the functional relevance of the findings. Comparing the observed protein changes in early and late DKD, consistent upregulation of 21 and downregulation of 18 proteins was detected. Among these were downregulated peroxisomal proteins such as NUDT19, ACOX1, and AMACR and upregulated mitochondrial proteins related to aminoacid metabolism including GLS, GLDC, and GCAT. Several of these changes were also observed in the kidney cortex proteome of db/db mice. IHC of human kidney further confirmed the differential expression of NUDT19, AGPS, AMACR and CAT proteins in DKD.

**Conclusions:** Our study shows an extensive differential expression of peroxisomal proteins in the early stages of DKD that persists regardless of the disease severity. These proteins therefore represent potential markers of early DKD pathogenesis. Collectively, essential pathways associated with peroxisomes such as lipid β-oxidation, plasmalogen synthesis, aminoacid metabolism and response to oxidative stress are downregulated in early DKD, providing new perspectives and potential markers of diabetic kidney dysfunction.

## INTRODUCTION

According to the 2019 Atlas of the International Diabetes Federation, about 463.0 million adults have type 1 (T1D) or type 2 diabetes (T2D) (1,2). Diabetic patients are at high risk of developing diabetic kidney disease (DKD), cardiovascular disease (CVD), neuropathy and retinopathy (1).

The prevalence of DKD is increasing and is associated with a heavy societal and financial burden (3). DKD is characterized by altered glomerular filtration and proteinuria resulting in up to 50% of the end stage kidney disease (ESKD) patients due to DKD (4,5). DKD is characterized by kidney ultra-structural and morphological alterations such as mesangial expansion, nodular glomerular sclerosis, glomerular basement membrane (GBM) thickening, and tubulointerstitial fibrosis (6). Glomerular injury characterizes the early stages of DKD, thus glomeruli are significant targets to investigate the molecular mechanisms of early DKD pathogenesis (7).

Several biological processes relevant to DKD have been studied such as mitochondrial dysfunction, reactive oxygen species (ROS) production (8,9), NADPH oxidase (NOX) activity, podocyte apoptosis and autophagy (9) leading to glomerular injury (10). Oxidative stress has been highlighted as a significant contributor to DKD (10) and progression to ESKD (11), being directly linked to podocyte damage, proteinuria, and tubulointerstitial fibrosis (11). Oxidative stress is triggered from changes in kidney lipid metabolism (12) with kidney lipotoxicity and lipid accumulation being considered pathological hallmarks of DKD (12,13). Glomerular lipid accumulation could lead to podocyte death and insulin resistance (12,13). Despite this accumulated knowledge, the absence of efficient inhibitors for the progressive distortion of kidney structure and function reflects the gaps in our understanding of DKD pathogenesis (14,15). The most promising treatment of DKD currently is inhibition of sodium-glucose transporter 2 (SGLT2). SGLT2 inhibitors were initially developed to lower blood glucose concentrations, but also showed very favorable protective effects in DKD apparently independent of the blood glucose lowering effect (15).

Many mouse and rat models of T1D and T2D have been established in order to dissect diabetic and DKD pathogenesis (16). Several DKD studies use Ins2Akita mice as model of T1D since these mice have glomerular basement membrane thickening, increased albumin excretion, glomerulosclerosis and interstitial fibrosis, mimicking human DKD (17). Hyperglycemia in Ins2Akita mice is thought to induce oxidative stress, resulting in kidney injury (18). Also the db/db mouse model is frequently used as a model of human T2D due to susceptibility to obesity, insulin resistance and T2D resulting from leptin deficiency, and development of progressive histological lesions in their kidneys (16).

To shed more light onto the molecular mechanisms of early DKD pathogenesis and progression, our study targeted the comprehensive molecular characterization of kidney tissue compartments at different developmental time points from the widely used Ins2Akita model of DKD with subsequent validation of findings in the db/db model and in human kidney biopsies. We performed high-resolution, quantitative mass spectrometry (MS)-based proteomics analysis of kidney glomeruli or cortex from the mouse models. Our results highlight a conserved (throughout T1D and T2D models and humans) downregulation of peroxisomal function and their cross-talk with mitochondria in early and late DKD, opening up new avenues for DKD therapy.

## METHODS

### Animals

The mouse models C57BL/6-Ins2Akita/J (Ins2Akita-T1D) and BKS.Cg-+Leprdb/+Leprdb/OlaHsd (dbdb-T2D) were used. In the glomerular proteome study 4 groups of animals were included, 2-month-old Ins2Akita (INS2) (n=8) and respective controls (WT2; n=7); and 4-month-old Ins2Akita (INS4) (n=8) and respective controls (WT4, n=8) (7). For the kidney cortex proteome study db/db mice were used. The former included 6-month-old db/db (n=3) and respective controls db/dm: (n=5).

### Isolation of glomeruli

Glomeruli were isolated as previously described in (7) Klein J, et al. 2020. In brief, mice were anesthetized and a catheter was placed into the abdominal aorta. The lower part of the abdominal aorta was perfused with 40 mL of Dynabeads M-450 Tosylactivated (4.5 µm diameter, Dynal A.S., Oslo, Norway) at a concentration of 2 × 10^6^ beads/mL followed by 15 mL of cold PBS. This procedure allows accumulation of beads in glomeruli. Next, the left and right kidneys were collected and decapsulated.

The left kidney was gently pressed manually through a 70 µm cell strainer using a flattened pestle followed by washing of the cell strainer with 20 mL of cold PBS. The filtrate was centrifuged at 200×*g* for 5 min at 4 °C, and the glomerular pellet was adjusted to 2 mL with PBS and transferred to an Eppendorf tube that was placed in a magnetic particle concentrator (Dynal A.S., Oslo, Norway) to concentrate the glomeruli into a pellet. The supernatant was discarded and the pellet was washed 5 × with 1 mL of PBS. The final pellet was resuspended in 100 µL PBS. This procedure allows the isolation of ∼ 4,000 glomeruli per kidney.

Based on light microscopy survey, our glomerular suspensions were highly enriched for glomeruli. In addition, mRNA quantification showed that isolated glomeruli were enriched in glomerular-specific genes.

### Murine kidney histology

Kidney histology was performed as described in (7) Klein J, et al. 2020. In brief, kidney lesions were assessed by specific (immuno) histological evaluation of the kidney structure (PAS) and fibrosis (glomerulo and tubulointerstitial fibrosis, Masson-trichrome and Sirius red staining, collagen III staining). At least 50 glomeruli including superficial and juxtaglomerular cortical area, were examined for each animal.

### Biochemical analysis

Biochemical analysis was performed as described in (7) Klein J, et al. 2020. Urinary albumin concentration was measured by ELISA using the AlbuWell kit (WAK-Chemie Medical GmbH, Steinbach, Germany). Urinary creatinine concentration was measured by the colorimetric method of Jaffe. Blood glucose levels were measured in caudal blood using a glucometer.

### Sample preparation for proteomics

Sample preparation was performed as described in Latosinska A, et.al. 2020 (19). In brief, tissue samples were homogenized in lysis buffer consisting of 7M Urea, 2M Thiourea, 4% CHAPS and 1% DTE and processed with the GeLC-MS method (20). Ten micrograms of each sample were analyzed in SDS-PAGE (5% stacking, 12% separating). Electrophoresis was stopped when samples just entered the separating gel. Gels were fixed with 30% methanol, 10% acetic acid for 30 min followed by 3 washes with water (5 min each) and stained with Coomassie Colloidal Blue overnight. Excess of stain was washed with water. Each band was excised from the gel and further sliced into small pieces (1-2 mm). Gel pieces were destained with 40% Acetonitrile, 50 mM NH4HCO3 and then reduced with 10 mM DTE in 100 mM NH4HCO3 for 20 min RT. After reduction, samples were alkylated with 54 mM Iodoacetamide in 100 mM NH4HCO3 for 20 min at room temperature in the dark. Samples were then washed with 100 mM NH4HCO3 for 20 min at room temperature, followed by another wash with 40% Acetonitrile, 50 mM NH4HCO3 for 20 min at room temperature and a final wash with ultrapure water under the same conditions was performed. Gel pieces were dried in a centrifugal vacuum concentrator and trypsinised overnight in the dark at room temperature, by adding 600 ng of trypsin per sample (trypsin stock solution: 10 ng / μL in 10 mM NH4HCO3, pH 8.5). Peptides were extracted after incubation with the following buffers: 50 mM NH4HCO3 for 15 min at room temperature followed by two incubations with 10% Formic Acid, Acetonitrile (1:1) for 15 min at room temperature. Peptides were eluted in a final volume of 600 μL and filtered with 0.22 μm PVDF filters (Merck Millipore) before dried in a centrifugal vacuum concentrator. Dried peptides were reconstituted in mobile phase A buffer (0.1% formic acid, pH 3.5) and processed with LC-MS/MS analysis.

### LC-MS/MS analysis

LC-MS/MS experiments were performed on the Dionex Ultimate 3000 UHPLC system coupled with the high resolution nano-ESI Orbitrap-Elite mass spectrometer (Thermo Scientific). Each sample was reconstituted in 10 μL loading solution composed of 0.1 % v/v formic acid. A 5 μL volume was injected and loaded on the Acclaim PepMap 100, 100 μm × 2 cm C18, 5 μm, 100 Ȧ trapping column with the ulPickUp Injection mode with the loading pump operating at flow rate 5 μL/min. For the peptide separation the Acclaim PepMap RSLC, 75 μm × 50 cm, nanoViper, C18, 2 μm, 100 Ȧ column retrofitted to a PicoTip emitter was used for multi-step gradient elution. Mobile phase (A) was composed of 0.1 % formic acid and mobile phase (B) was composed of 100% acetonitrile, 0.1% formic acid. The peptides were eluted under a 240 minutes gradient from 2% (B) to 80% (B). Flow rate was 300 nL/min and column temperature was set at 35 °C. Gaseous phase transition of the separated peptides was achieved with positive ion electrospray ionization applying a voltage of 2.5 kV. For every MS survey scan, the top 10 most abundant multiply charged precursor ions between m/z ratio 300 and 2200 and intensity threshold 500 counts were selected with FT mass resolution of 60.000 and subjected to HCD fragmentation. Tandem mass spectra were acquired with FT resolution of 15.000. Normalized collision energy was set to 33 and already targeted precursors were dynamically excluded for further isolation and activation for 45 sec with 5 ppm mass tolerance.

### MS data processing

Raw files were analyzed with Proteome Discoverer 1.4 software package (Thermo Finnigan), using the SEQUEST search engine and the UniProt mouse (Mus musculus) reviewed database, downloaded on November 22, 2019 including 16.935 entries and processed for protein identification and relative quantification under stringent criteria by the use of well-established pipelines in our lab (21). The search was performed using carbamidomethylation of cysteine as static and oxidation of methionine as dynamic modifications. Two missed cleavage sites, a precursor mass tolerance of 10 ppm and fragment mass tolerance of 0.05 Da were allowed. False discovery rate (FDR) validation was based on q value: target FDR: 0.01. Label free quantification was performed by utilizing the precursor ion area values exported from the total ion chromatogram as defined by the Proteome Discoverer 1.4 software package.

Output files from Proteome Discoverer were processed with an in-house script in the R environment for statistical computing (version 4.0.3) as follows: Protein lists were concatenated into a master table. Raw protein intensities for each individual sample were subjected to normalization according to X’ = X/Sum(Xi) * 10^6 and only proteins present in at least 55% of the samples in at least one group were further selected for downstream statistical analysis.

### Statistical Analysis of proteomics data

Statistical significance of continuous variables was defined at p < 0.05 with the non-parametric Mann-Whitney test. Proteins with p value ≤0.05 and ratio ≥1.5 (upregulated) or ≤0.67 (downregulated) were considered statistically significant differentially expressed. Dotplots for the mouse physiopathologic characterization were created with functionality from the ggplot2 and ggpubr R packages. Depicted statistical comparisons correspond to independent Mann-Whitney tests. For the correlation analysis, Spearman’s correlation was utilized to assess relationships of our data to one external proteomics dataset (22), using the means of the normalized protein intensity across samples after transforming them to the natural logarithmic scale. Heatmap was created with the gplot package, after Z-scaling of the normalized protein intensities. Euclidean distance and Ward’s hierarchical method (option: ward.D2) were selected for the clustering of both rows and columns. Graphing and statistical analysis were performed in the RStudio environment (R version 4.0.3).

### Functional analysis

Pathway annotation was performed with the ClueGO plug-in 3.7.2 of Cytoscape platform using REACTOME pathways database (updated on February 17, 2020). Biological function annotation was performed using the ClueGO plug-in 3.7.2 of Cytoscape platform using GO-Biological Process EBI-Uniprot GOA database (updated on February 17, 2020). Only statistical significant pathways (Benjamini-Hochberg (BH) corrected p-value ≤ 0.05, two-sided hypergeometric test) were taken into account. For the remaining parameters, default settings were used. Results were simplified based on biological relevance and only the leading term from each group is presented.

### Investigation through transcriptomics data analysis

Nephroseq (www.nephroseq.org,) was employed for the investigation of the expression of the shortlisted mitochondrial and peroxisomal proteins in existing mouse and human transcriptomics datasets. The list of proteins was uploaded in Nephroseq v4 in the form of EntrezGene IDs. DKD datasets selection was held after application of the filters: Primary Filters > Group > Diabetic nephropathy. The corresponding gene expression was searched in 7 available DKD mouse and human datasets observed after filtering, on comparison of DKD vs. Healthy Living Donor groups (TableS1). Only significantly deregulated genes (p < 0.05) were extracted and their differential expression was compared with the mitochondrial and peroxisomal deregulated proteins.

### Clinical material

Within a period of 8 years (2013 – 2020) a total number of 100 kidney biopsies diagnosed as DKD were retrieved from the human Renal Biopsies archive of the 1st Department of Pathology of Athens (National and Kapodistrian University of Athens, Medical School). All cases were initially classified based on their glomerular lesions into the 4 classes of DKD (I, IIa/b, III and IV) according to Tervaert et al. (23). Interstitial fibrosis and tubular atrophy (IFTA) as well as interstitial inflammatory infiltration and vascular lesions (arteriolar hyalinosis, arteriosclerosis) were also studied and scored from 0 to 9 according to Tervaert et al. (23) in order to assess the severity and chronicity of DKD.

Information on gender, age, serum-creatinine value and albuminuria levels was collected and the estimated Glomerular Filtration Rate (eGFR) was calculated using the online National Kidney Foundation GFR Calculator (24). Combining eGFR and albuminuria levels the cases were classified to 5 clinical stages (G1, G2, G3a/b, G4 and G5 – A1, A2 and A3) of chronic kidney disease (CKD) due to diabetes mellitus (DM) according to KDIGO Guidelines (25). All cases were classified as A3 since albuminuria levels were always >0.3g/24h.

Sixteen out of the 100 cases were selected in order to assess the immunohistochemical expression of the peroxisomal proteins Nucleoside diphosphate-linked moiety X motif 19 (NUDT19), Alpha-methylacyl-CoA racemase (AMACR), Peroxisomal catalase (CAT) and Alkyldihydroxyacetonephosphate synthase (AGPS) in human renal tissues with DKD.

The selected cases met the following criteria: 1) Absence of coexisting non diabetic kidney disease, 2) Adequate representation of the histopathological lesions of DKD, reflecting the 4 histological classes of DKD and 3) Kidney function deterioration, as it is reflected in CKD stage, equivalent to DKD lesion progression.

Additionally, normal kidney tissue showing no signs of DKD, CKD or other kidney pathology, obtained from radical nephrectomy specimens were used as controls.

The main clinicopathological parameters of the 16 cases are shown in TableS2.

### Immunohistochemistry of human DKD specimens

Immunohistochemical staining for the markers under study was performed on 4-μm-thick formalin-fixed paraffin sections. After heating overnight at 37°C, deparaffinization, rehydration and antigen retrieval were performed in one step, by treating the slides in an automated module (PT Link, Dako) for 20 min at 96°C with the reagents EnVision FLEX Target Retrieval Solution High pH (50x) (Dako) for CAT and EnVision FLEX Target Retrieval Solution Low pH (50x) (Dako) for NUDT19, AGPS, AMACR. Endogenous peroxidase activity was quenched with 0.3% hydrogen peroxide in Tris-buffered saline (TBS), for 15 min. After rinsing with TBS, normal horse serum was applied for 20 min, to block non-specific antibody binding. Subsequently, sections were incubated overnight at 4°C with the primary antibodies as follow: Anti-AMACR (rabbit polyclonal) (Atlas Antibodies, Sweden) at a dilution 1:2000, anti-AGPS (rabbit polyclonal) (Atlas Antibodies, Sweden) at a dilution 1:50, anti-CAT (rabbit polyclonal) (Atlas Antibodies, Sweden) at a dilution 1:3000 and anti-NUDT19 (EPR13162-63) (rabbit monoclonal) (abcam, Cambridge, UK) at a dilution 1:100. A two-step technique (polymer, HRP-conjugated; Vector Laboratories, Burlingame, CA, USA) was used for visualization, with diaminobenzidine as a chromogen. Finally, sections were counterstained with haematoxylin and mounted.

### Evaluation of immunohistochemistry

Qualitative immunohistochemical evaluation was assessed by two independent observers (HG and DP) blinded to clinical data. The pattern, topography, extension, and intensity of staining of all the under study antibodies in both the controls and DKD cases were studied. A rough comparison of the expression of each marker between the controls and DKD cases, as well as between cases of different DKD classes was also performed.

Staining intensity was also further quantified using ImageJ software as previously described (20). Optical density was normalized over the unstained tissue and mean intensity values were estimated. Statistical significance among multiple groups (stages) was confirmed with ANOVA analysis and further pair-wise comparisons were performed with Student’s T-test. Values of p<0.05 were considered as statistically significant.

## RESULTS

### 1. Physiopathologic characterization of Ins2Akita and db/db mouse models

Our study was performed using the Ins2Akita mouse model representing early to late kidney morphological lesions and dysfunction as observed, respectively, in T1D DKD (15). Further, db/db mice were investigated, displaying insulin resistance, obesity, T2D and progressive kidney histological lesions as observed in human T2D (15). Ins2Akita mice became hyperglycemic at 2 month of age (Figure S1A) and exhibited significant increased urinary albumin to creatinine ratio (ACR) at 4 months (Figure S1A) compared to WT mice. Six month old db/db mice displayed hyperglycemia (Figure S1B) and increased ACR compared to control mice (db/dm) (Figure S1B).

### 2. Glomerular proteome profiles from Ins2Akita mice reveal prominent changes in mitochondrial and peroxisomal proteins in early and late DKD

Proteomics analysis was applied to investigate the differences in the molecular profiles of glomeruli between Ins2Akita mice 2 month (INS2) (early DKD) and 4 month (INS4) (late DKD) old and respective controls (Table 1). An average of 1550 proteins in wild type mice and 1600 proteins in Ins2Akita mice were detected (TableS3). The full list of identified proteins using a 55% frequency threshold is shown in TableS4-Sheet1 (Ins2Akita glomerular dataset).

**Table 1.**
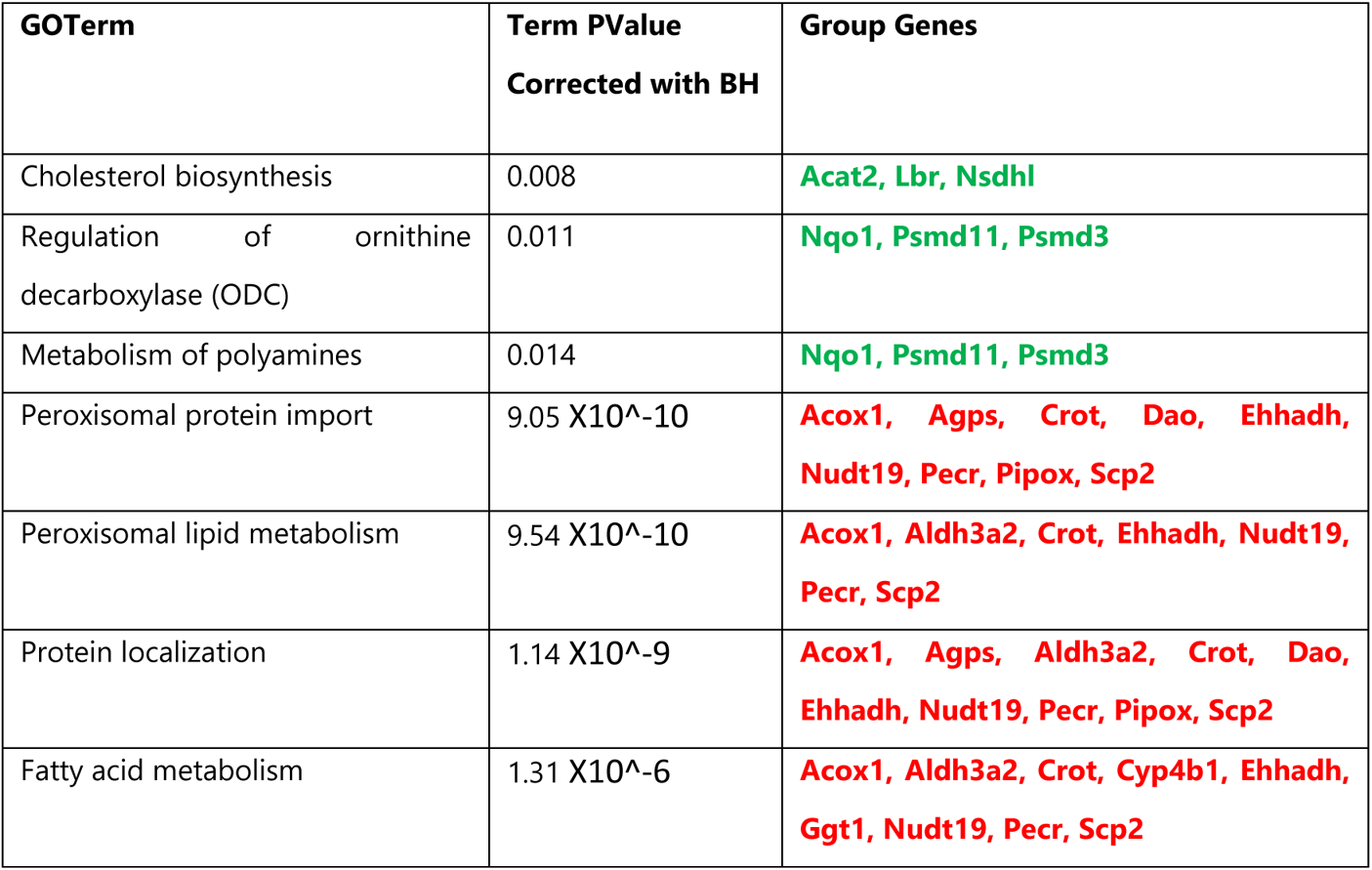
Significant pathways represented by glomerular proteins associated with early DKD according to the BH corrected p value of the GO Term. The upregulated (green color) and downregulated proteins (red color) in Ins2Akita compared to WT controls are shown below.

These protein lists largely overlapped with similar published proteomics datasets (Waanders LF, et al. 2009 (22)). In brief, most of the identified proteins in our study were also detected in these published data (60% for wild type mice and 73% for the Ins2Akita) (22). Correlation analysis of the protein abundance detected in our study with Waanders LF, et al. 2009 (22) resulted in correlation factor R= 0.7 for both, the wild type and the Ins2Akita mice (Figure S2), demonstrating consistency between the different studies.

Multiple pair-wise comparisons were performed to identify the consistent changes associated with early DKD (TableS4-Sheet1).

When comparing the abundance of proteins of 2 month old Ins2Akita to the respective 2 month WT controls, 100 proteins were significantly up-regulated (TableS4-Sheet2) and 77 were down-regulated (TableS4-Sheet3) in Ins2Akita. Similarly, comparison of the abundance of proteins between Ins2Akita and WT 4 month old mice revealed 155 proteins with significantly increased abundance (TableS4-Sheet2) and 111 with decreased abundance in Ins2Akita (TableS4-Sheet3).

The biological functions represented by the deregulated glomerular proteins associated with early DKD are predominately related to cholesterol metabolic process, glycine metabolic process, lipid storage, cellular amino acid catabolic process, oxidoreductase activity, and fatty acid beta-oxidation (TableS5-Sheet1; respective pathways are summarized in Table 1). The most prominent biological functions of the proteins significantly changed in late DKD are branched-chain amino acid catabolic process, glutamate and glutamine metabolic process, lipid storage, carboxylic acid catabolic process, fatty acid beta-oxidation, acyl-CoA metabolic process, and oxidoreductase activity (TableS5-Sheet2; a summary of main pathways is provided in Table 2).

**Table 2.**
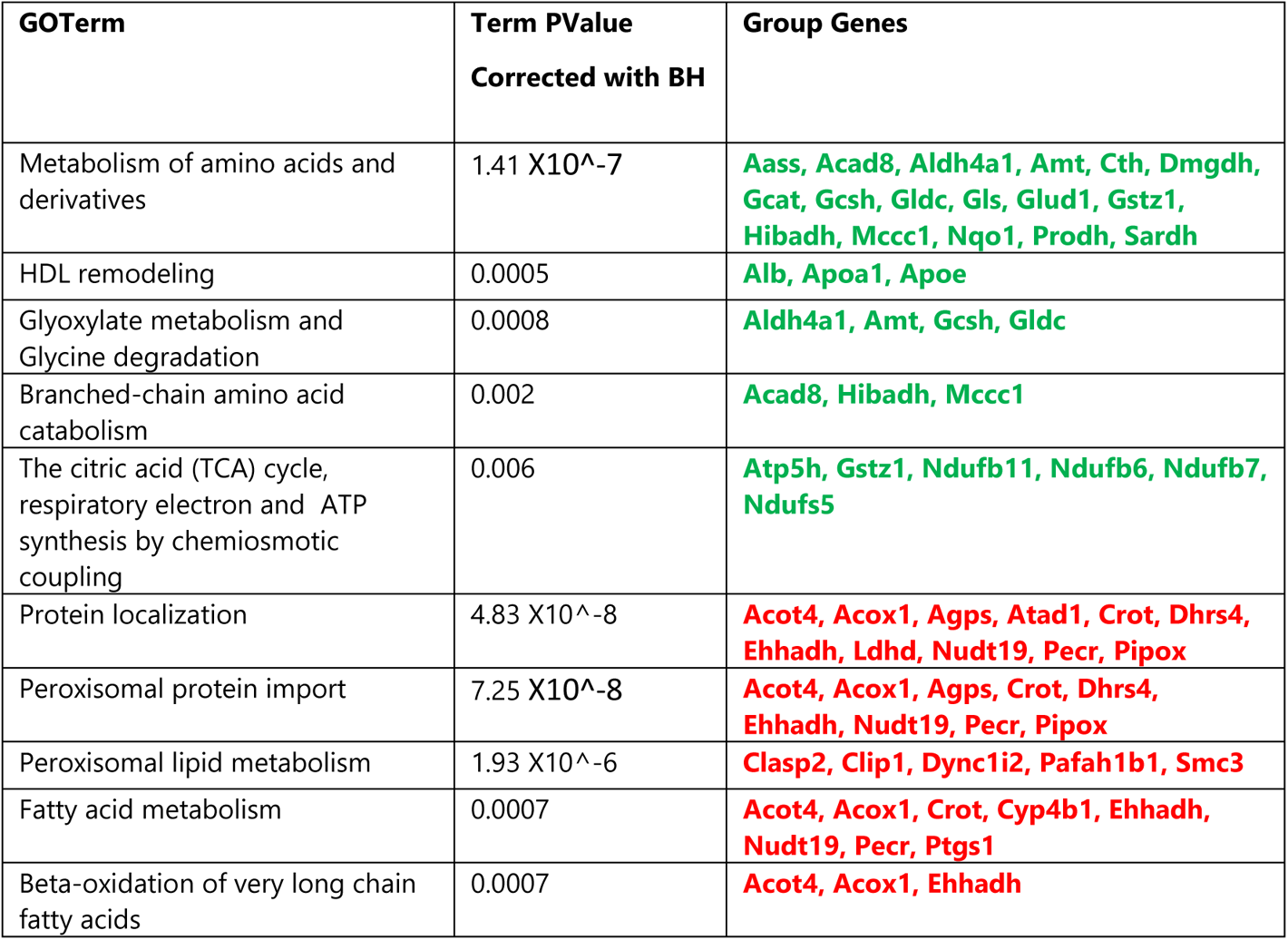
Significant pathways represented by glomerular proteins associated with late DKD according to BH corrected p value of the GO Term. The upregulated (green color) and downregulated proteins (red color) in Ins2Akita compared to WT controls are shown below.

To verify the observations and identify early changes associated with the disease, glomerular proteins that prominently changed in diabetic animals in early and late stages were studied further. Twenty-five upregulated and 18 downregulated proteins with consistently changed trend between the stages of DKD were selected (TableS4-Sheet4).

To correct for any potential impact of aging, the differences from the comparison WT4 VS WT2 (TableS6) were subtracted from the above glomerular proteins associated with DKD; this gave rise to a final list of 39 proteins (21 up and 18 down-regulated) with consistent expression trend (increased / decreased abundance) in early and late - DKD versus controls (Figure 1; Table 3 and Table 4, respectively).

**Figure 1.**
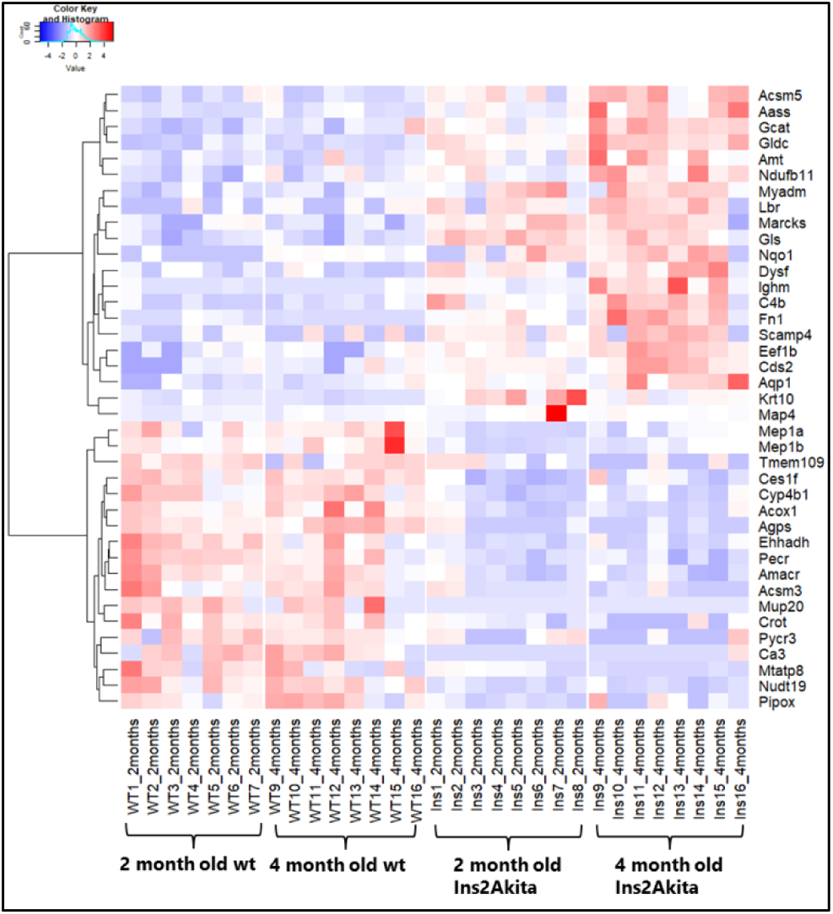
The expression changes of the 39 (21 up and 18 down-regulated) in early and late DKD versus controls are illustrated in the heatmap for each group, indicating the similar trend of expression of these proteins in the two disease stages.

**Table 3.** Twenty-one consistently upregulated glomerular proteins in early and late DKD versus controls. Of these proteins, 7 are related to diabetes (the references are presented in TableS7) and 15 could be attributed to kidney expression and/or function (the references are presented in TableS7).

| **Accession** | **Name** | **pval_INS2**<br>**vsWT2** | **Ratio_INS2**<br>**vsWT2** | **pval_INS4**<br>**vsWT4** | **Ratio_INS4**<br>**vsWT4** | **Related**<br>**to**<br>**Diabetes** | **Kidney**<br>**expression**<br>**and/or**<br>**function** |
| --- | --- | --- | --- | --- | --- | --- | --- |
| D3Z7P3 | Glutaminase kidney isoform, mitochondrial OS | 0.000311 | 2.94 | 0.000311 | 2.33 | YES | YES |
| Q91W43 | Glycine dehydrogenase (decarboxylating), mitochondrial | 0.000311 | 2.16 | 0.000155 | 2.88 | YES | YES |
| P02535 | Keratin, type I cytoskeletal 10 | 0.00124 | 4.12 | 0.000155 | 3.62 | NO | NO |
| O88986 | 2-amino-3-ketobutyrate coenzyme A ligase, mitochondrial | 0.00214 | 3.54 | 0.00295 | 2.57 | YES | YES |
| Q99K67 | Alpha-aminoadipic semialdehyde synthase, | 0.00218 | 1.76 | 0.00295 | 2.57 | YES | YES |
| P26645 | Myristoylated alanine-rich C-kinase substrate | 0.00314 | 2.21 | 0.00986 | 2.58 | NO | YES |
| P01029 | Complement C4-B | 0.00897 | 5.57 | 0.00295 | 4.42 | NO | NO |
| Q99L43 | Phosphatidate cytidyltransferase 2 | 0.0125 | 2.12 | 0.00295 | 1.92 | NO | YES |
| P27546 | Microtubule-associated protein 4 | 0.0128 | 6.06 | 0.0148 | 1.73 | NO | YES |
| Q02013 | Aquaporin-1 | 0.0128 | 2.23 | 0.00187 | 2.24 | NO | YES |
| Q9JKV5 | Secretory carrier-associated membrane protein 4 | 0.0173 | 2.065 | 0.033 | 2.59 | NO | YES |
| P01872 | Ig mu chain C region | 0.0231 | 4.94 | 0.00147 | 14.0 | NO | NO |
| Q3U9G9 | Lamin-B receptor | 0.0231 | 3.59 | 0.00817 | 2.284 | NO | NO |
| O09111 | NADH dehydrogenase [ubiquinone] 1 beta | 0.0289 | 1.898 | 0.00699 | 1.94 | YES |  |
| Q8BGA8 | Acyl-coenzyme A synthetase ACSM5, mitochondrial | 0.0289 | 1.52 | 0.000622 | 1.95 | NO | YES |
| P11276 | Fibronectin | 0.0292 | 3.05 | 0.000554 | 103 | YES | YES |
| O35682 | Myeloid-associated differentiation marker | 0.0321 | 2.85 | 0.00699 | 2.25 | NO | NO |
| O70251 | Elongation factor 1-beta | 0.0321 | 1.76 | 0.00135 | 2.30 | NO | YES |
| Q9ESD7 | Dysferlin | 0.0321 | 2.16 | 0.00377 | 4.55 | NO | YES |
| Q8CFA2 | Aminomethyltransferase,<br>mitochondrial | 0.04 | 2.17 | 0.00295 | 2.95 | YES | YES |
| Q64669 | NAD(P)H dehydrogenase<br>[quinone] 1 | 0.043 | 6.96 | 0.01 | 1.60 | NO | YES |

**Table 4.**
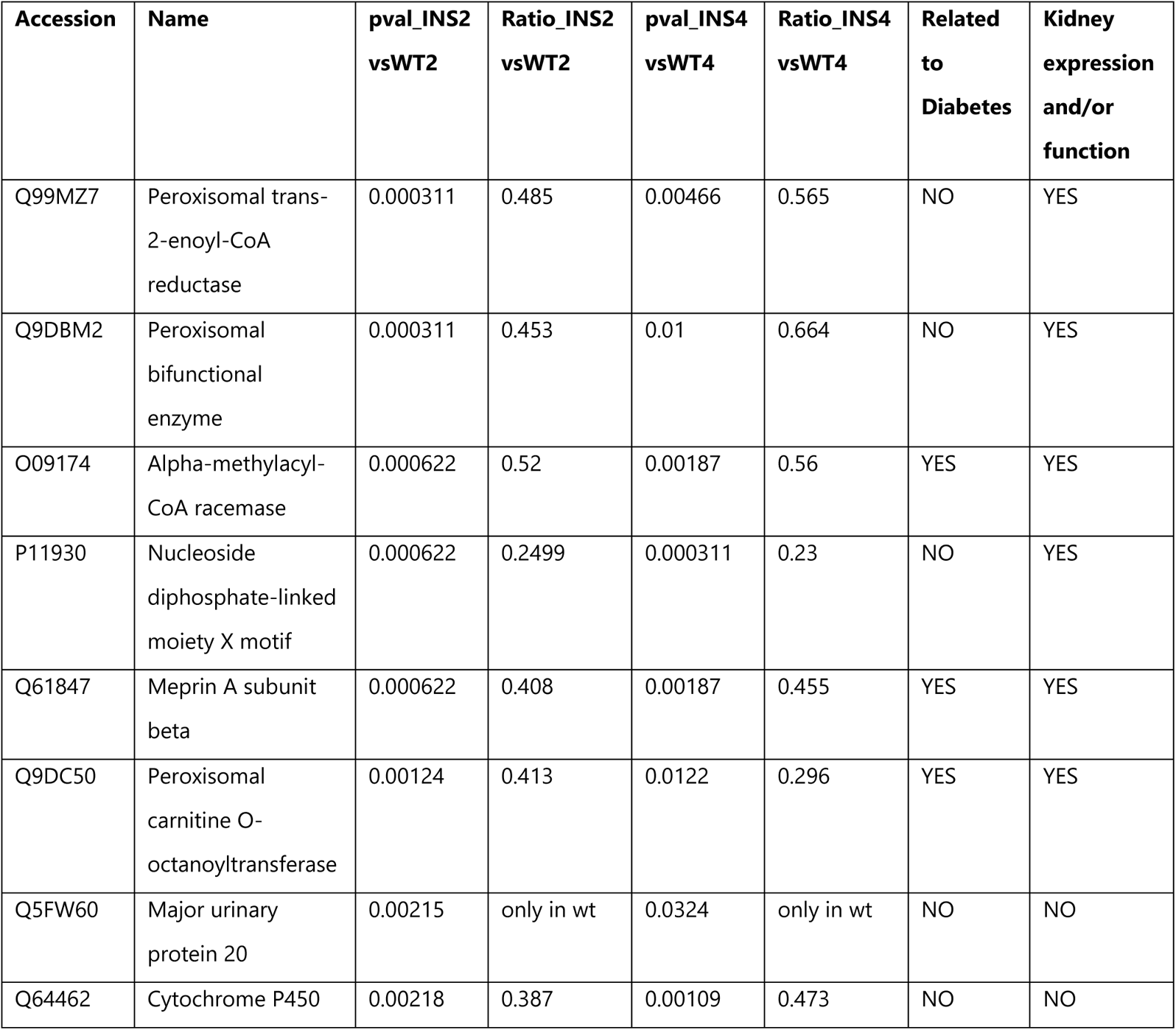
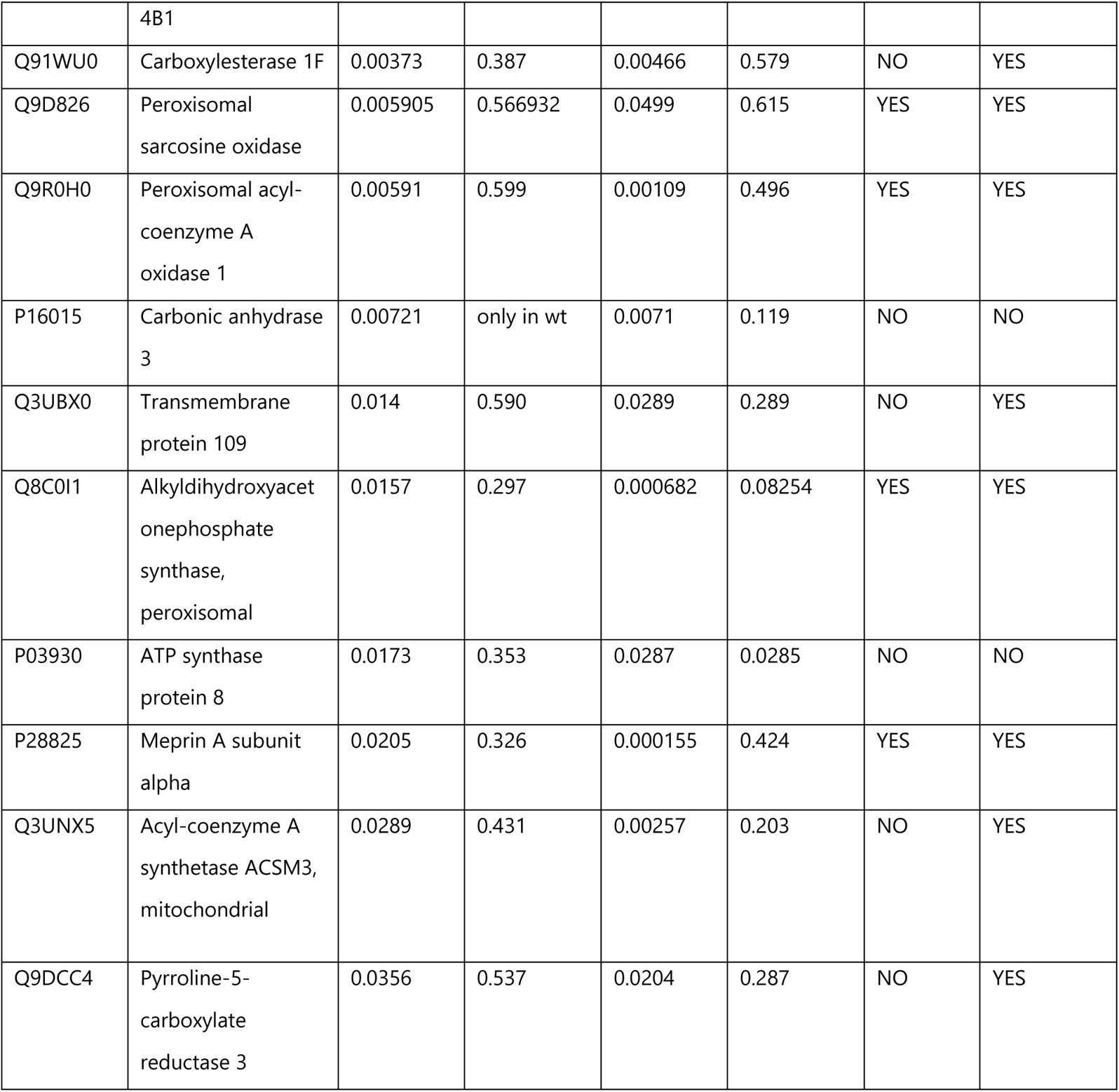
Eighteen consistently downregulated glomerular proteins in early and late DKD. Of these proteins, 7 are related to diabetes (the references are presented in TableS7) and 14 could be attributed to kidney expression and/or function (the references are presented in TableS7).

According to the literature and the databases Uniprot and Human Protein Atlas, among the 21 upregulated proteins, 7 are already related to diabetes (Table 3 and in detail in TableS7) and 15 have attributed kidney expression and/or function (Table 3 and in detail in TableS7). Among the 18 downregulated proteins, 7 are related to diabetes (Table 4 and in detail inTableS7) and 14 have documented kidney expression and/or function (Table 4 and in detail in TableS7).

The biological functions of the consistently changed glomerular proteins associated with early and late DKD are listed in Table 5.

**Table 5.** The biological function of the consistently changed glomerular proteins associated with early and late DKD. The upregulated (green color) and downregulated proteins (red color) in Ins2Akita compared to WT controls are shown below.

| **GO Term** | **Group P Value Corrected with BH** | **Group Genes** |
| --- | --- | --- |
| cellular amino acid catabolic process | 2.51 X10<sup>-6</sup> | **Aass,Amt,Gcat,Gldc,Gls** |
| alpha-amino acid catabolic process | 2.83 X10<sup>-6</sup> | **Aass,Amt,Gcat,Gldc,Gls** |
| response to mercury ion | 0.0004 | **Aqp1,Cds2,Dysf,Fn1,Krt10** |
| cellular response to osmotic stress | 0.00208 | **Aqp1,Cds2,Dysf** |
| fatty acid catabolic process | 2.36 X10<sup>-7</sup> | **Acox1,Ces1f,Crot,Ehhadh,Pipox,Pycrl** |
| isoprenoid catabolic process | 0.0218 | **Amacr** |
| ether lipid biosynthetic process | 0.0275 | **Agps** |
| trans-2-enoyl-CoA reductase (NADPH) activity | 0.029 | **Pecr** |

As shown in Table 5, many of the upregulated proteins are involved in aminoacid metabolism performed in mitochondria whereas the downregulated proteins are mainly involved in fatty acid catabolism and ether lipid biosynthesis which are recognized as basic peroxisomal functions.

### 3. Validation of findings in late T2D DKD animal model and cross-omics validation, further confirming peroxisomal changes in DKD

In order to validate the findings of the glomerular proteome analysis and investigate further if they are observed also in a T2D DKD model, the kidney cortex proteomic profile of db/db mice, a model of T2D DKD, of advanced age (6 months) was also analyzed. The diabetic stage (late DKD) of the db/db mouse model was verified via biochemical data analysis (Suplementary Figure 1). Multiple pair-wise comparisons were performed (described below).

On average 674 proteins were detected in wild type and 670 in db/db mice (TableS8). A comprehensive list of the identified proteins including the fold change (ratio) and the statistics is shown TableS9-Sheet1 (db/db kidney cortex dataset).

When comparing the db/db to respective WT data, 33 upregulated proteins (TableS9-Sheet2), the majority related to glutamine and glutamate metabolic process and 23 downregulated proteins (TableS9-Sheet3), mostly related to peroxisomal protein import were identified. Interestingly, these processes were also observed in early T1D DKD (section 2). Taken together, these observations suggested that there exist common deregulated processes between early T1D, late T1D, and T2D DKD.

Several of the proteins associated with DKD identified in glomeruli from T1D mice were also found in the kidney cortex from T2D mice (Table 6). Specifically, NUDT19, AMACR and PIPOX, were downregulated in DKD in both glomeruli and kidney cortex in T1D and T2D models respectively (Table 6).

**Table 6.** Differentially expressed proteins with consistent expression trend in glomeruli and cortex collected respectively from Ins2Akita and db/db mice at different ages (months 2, 4, 6). The downregulated proteins are presented in red color.

| Description | Symbol | Ratio |  |  | Transcriptomics expression (Nephroseq; in DKD vs. controls) (ref) | Single-cell human kidney transcriptomics expression (Wilson et al. 29) |
| --- | --- | --- | --- | --- | --- | --- |
|  |  | db/db vs WT kidney cortex 6 months | Ins2Akita vs WT glomeruli 2 months | Ins2Akita vs WT glomeruli 4 months |  |  |
| Nucleoside diphosphate-linked moiety X | **NUDT19** | 0.236 | 0.25 | 0.23 | decrease (28) | not detected |
| Peroxisomal sarcosine oxidase | **PIPOX** | 0.406 | 0.567 | 0.615 | decrease (27) | not detected |
| Alpha-methylacyl-CoA racemase | **AMACR** | 0.185 | 0.52 | 0.56 | decrease (27) | decrease |

To further place our findings of deregulated mitochondrial and peroxisomal proteins in the context of published datasets in DKD, kidney transcriptomics data from DKD patients vs. healthy controls and DKD mouse versus healthy controls were retrieved from the Nephroseq database (7 datasets in total; 26–28). In addition, a reported single cell kidney transcriptomic dataset of early human DKD (29) was also investigated.

Of the reported differentially expressed and ‘consistent’ (Table 6) mitochondrial and peroxisomal proteins, mRNA expression levels were recorded for the 3 peroxisomal proteins NUDT19, PIPOX and AMACR with a reported decrease in DKD cases versus controls (Table 6).

The cross-omics analysis highlighted additional glomerular-cortical mitochondrial and peroxisomal proteins (Tables 3,4, TableS10, 11) in agreement in their change in mRNA and protein levels. In the transcriptomics datasets mRNA expression levels were also recorded for 3 mitochondrial proteins (GLS, GLDC, AMT) and 5 peroxisomal proteins (ACOX1, CROT, EHHADH, AGPS, and PECR), all consistently changing in early and late DKD (Tables 3,4) and 3 peroxisomal (CAT, EPHX2, DAO) cortical proteins associated with late DKD. For all these, with the exception of

AMT, changes at the mRNA levels in DKD vs healthy controls were in agreement to the observed changes in protein levels in our animal model experiments (TableS10, 11).

### 4. IHC validation of peroxisomal deregulation in human DKD patients

Given the consistency of these findings, a set of tissue sections of human kidney tissues was analyzed for the expression of the peroxisomal proteins NUDT19, AMACR, AGPS. In addition, the protein CAT that was found downregulated in the cortex of late DKD animal model (db/db mice) (TableS10, 11) was investigated, as a control, since it is the most studied and best-characterized peroxisomal antioxidant enzyme in kidney disease (66).

In a detailed qualitative analysis, NUDT19 (Figure 2) in normal kidney tissue showed tubular staining with a cytoplasmic, either coarsely granular or fine granular or pale homogeneous pattern. NUDT19 was detected in all types of tubules. In glomeruli, staining was found mainly in parietal cells of the Bowman’s capsule and occasionally in podocytes and mesangial cells (Figure 2A). In diabetic patients NUDT19 staining was seen in the kidney tubules of all cases, however, reduced compared to controls and with a decrease - in terms of both intensity and extension - from histological class I to class IV. Moreover, staining seemed more prominent in non-proximal convoluted tubules. Although NUDT19 was found only occasionally in the podocytes of the controls, podocytic expression (especially surrounding Kimmelstiel-Wilson nodules) was more intense in advanced cases of DKD.

**Figure 2.**
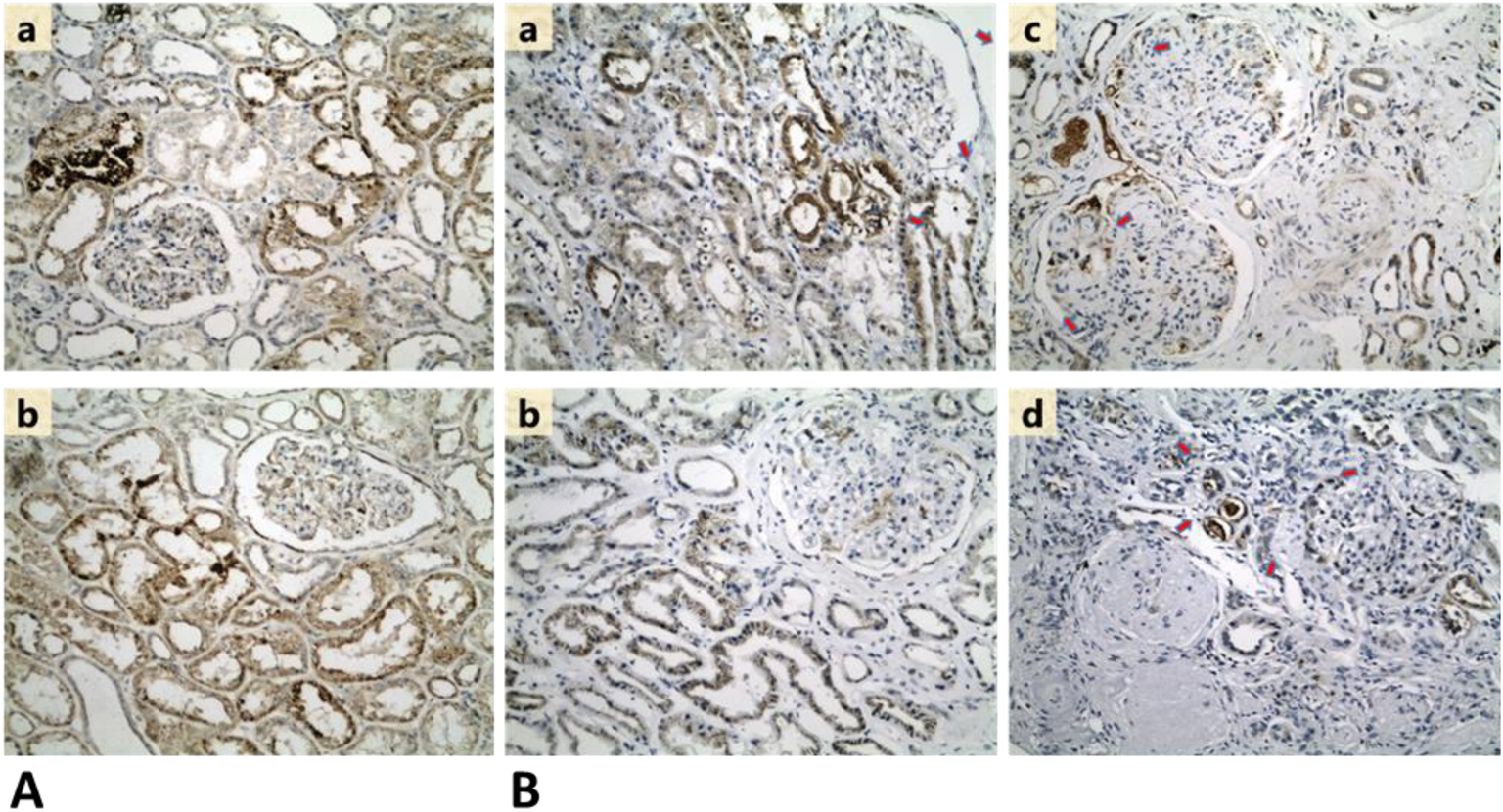
**A (a, b): Representative photos of NUDT19 staining in normal kidney tissue (x200).** Coarsely or fine granular and/or pale homogeneous cytoplasmic pattern of staining in all types of tubules and some glomerular cells. **B: Representative photos of NUDT19 staining in (a) class I, (b) class IIb, (c) class III and (d) class IV DKD (x200).** Gradual decrease from class I to IV can be seen. Note the intensification of podocytic staining around Kimmelstiel-Wilson nodules (red arrows).

AMACR in normal kidney tissue showed a diffuse, strong and densely granular, cytoplasmic staining in almost all proximal convoluted tubules (PCT), while distal convoluted tubules (DCT) demonstrated a focal weak granular cytoplasmic staining (Figure 3). Glomeruli were mostly negative, however some staining in parietal cells was also observed. Compared to the controls, DKD cases showed a significant decrease in AMACR staining intensity. AMACR staining also gradually decreased from histological class I to class IV DKD. Loss of AMACR expression was more often recorded as loss of staining in all the epithelial cells of any given tubule. However, in some tubules a segmental loss of expression was seen, with some epithelial cells retaining their cytoplasmic staining.

**Figure 3.**
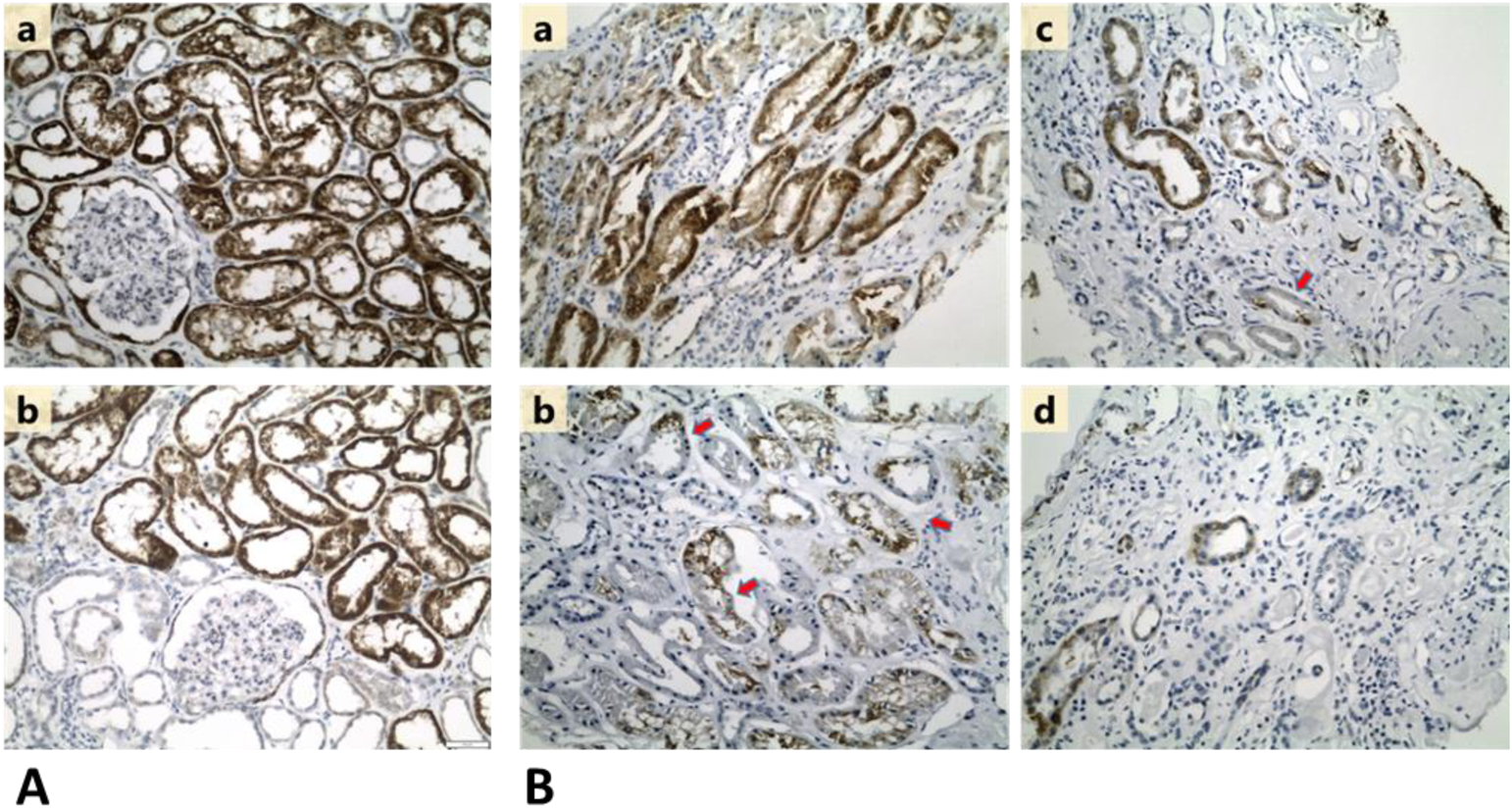
**A (a, b): Representative photos of AMACR staining in normal kidney tissue (x200).** Cytoplasmic staining, diffuse and strong in PCTs vs focal and weak in DCTs. Positivity in some parietal cells is also observed. **B: Representative photos of AMACR staining in (a) class I, (b) class IIb, (c) class III and (d) class IV DKD (x200).** Gradual loss of expression, usually in all the cells of any given tubule is observed. However, areas of segmental loss of staining can also be noted (red arrows).

In normal kidney tissue, AGPS showed a diffuse staining of moderate intensity in most kidney tubules with a cytoplasmic, membranous and occasionally nuclear pattern (Figure 4). In glomeruli, focal staining in parietal cells was seen. DKD cases exhibited a definite loss of AGPS expression, irrespectively of the DKD histological class. Of note, nuclear staining remained in some tubular epithelial cell nuclei.

**Figure 4.**
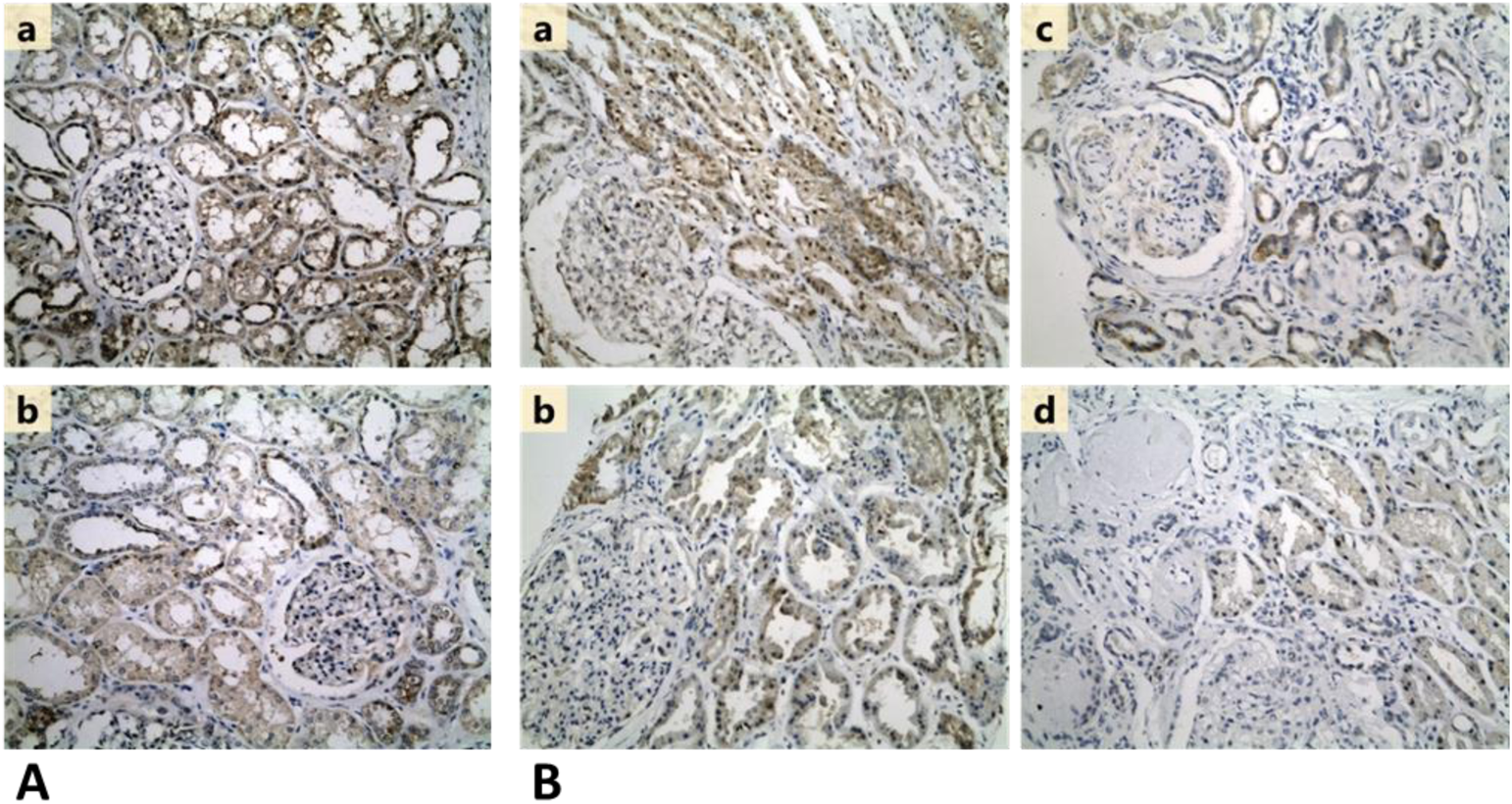
**A (a, b): Representative photos of AGPS staining in normal kidney tissue (x200).** Diffuse staining of moderate intensity with a cytoplasmic, membranous and occasionally nuclear pattern in most tubules and a few parietal cells. **B: Representative photos of AGPS staining in (a) class I, (b) class IIb, (c) class III and (d) class IV DKD (x200).** Decrease of expression compared to controls. Occasional nuclear pattern can still be noted in some cases.

Controls showed diffuse CAT staining in all types of renal tubules with a cytoplasmic, partly finely granular pattern of moderate intensity (Figure 5). In glomeruli, CAT staining was restricted to parietal epithelial cells. No endothelial cell staining was seen in vessels or glomerular capillaries. DKD cases demonstrated a definite loss of CAT expression, in comparison with the controls. Interestingly, in glomeruli of DKD cases, CAT was detected in the endothelial cells of the capillary walls, suggesting a possible endothelial activation under pathological conditions. However, no endothelial cell expression in extraglomerular vessels was seen. Another observation was a preferential CAT expression in tubules surrounding globally sclerosed glomeruli.

**Figure 5.**
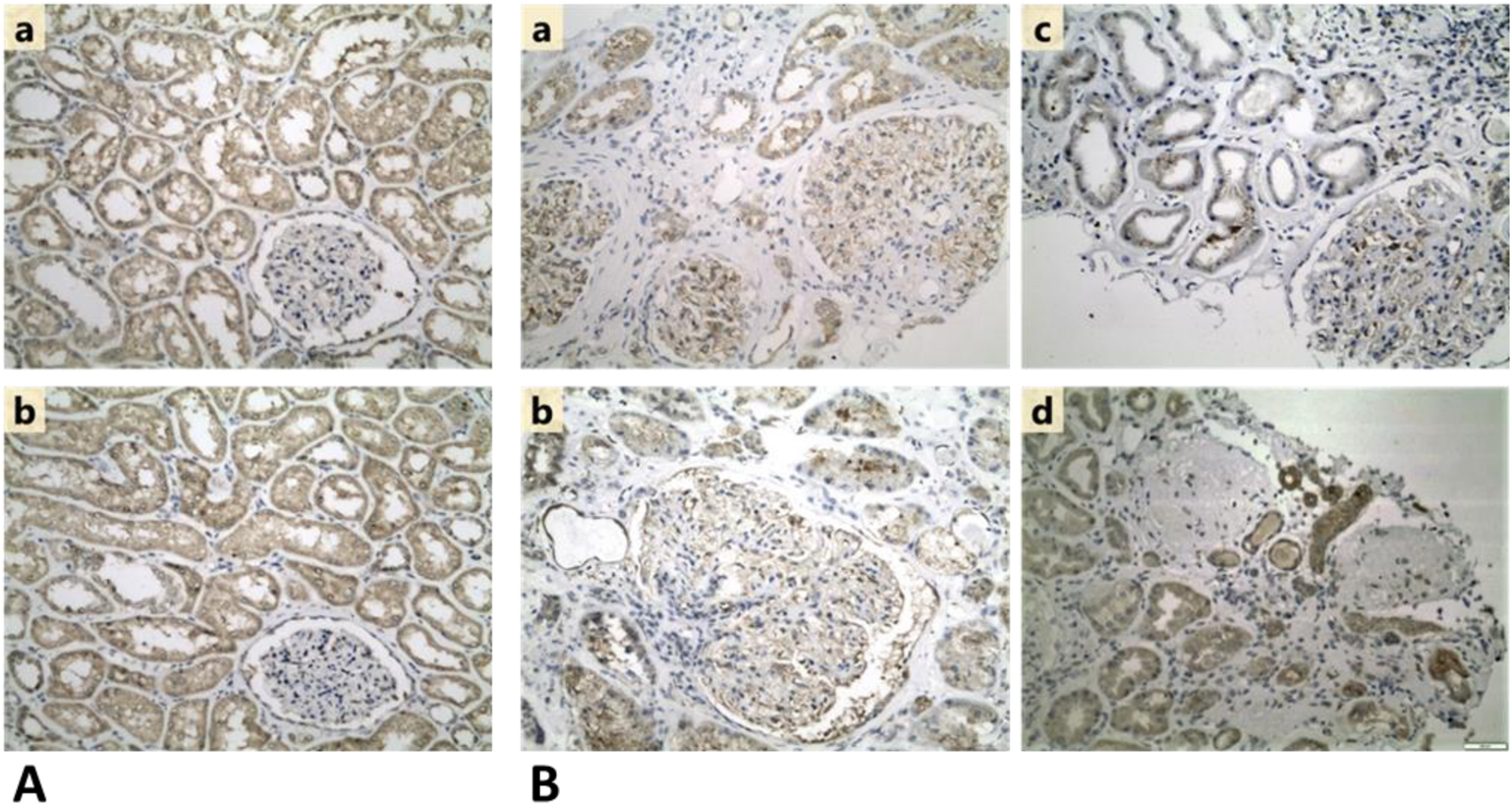
**A (a, b): Representative photos of CAT staining in normal kidney tissue (x200).** Diffuse,cytoplasmic staining of moderate intensity in tubules and parietal cells. **B: Representative photos of CAT staining in (a) class I, (b) class IIb, (c) class III and (d) class IV DKD.** Loss of CAT expression compared to controls. CAT positivity in endothelial cells of the capillary walls is observed in some cases (a, b and c). Preferential CAT expression in tubules surrounding globally sclerosed glomeruli can be noted (d).

Statistical significance among different groups was confirmed with ANOVA analysis (p<0.05) for CAT (p=0.0002), AMACR (p=0.0005) and AGPS (p=0.05), whereas in the case of NUDT19, a p> 0.05 was received. The latter may be attributed to the lower sample size among different groups for NUDT19 compared to the other proteins. However, it appears that the intensity of the protein stain in IHC as shown in Figures 2–5 follows the trend of the proteomics results obtained from the DKD animal model samples. Comparison of each group of samples for CAT, AMACR, AGPS and NUDT19 applying T-Test showed statistically significant difference among groups I+IIa compared to group IV for each of the aforementioned proteins (Figure 6). A control group (No DKD) was not included in the statistical analysis due to its small sample size (N=1).

**Figure 6.**
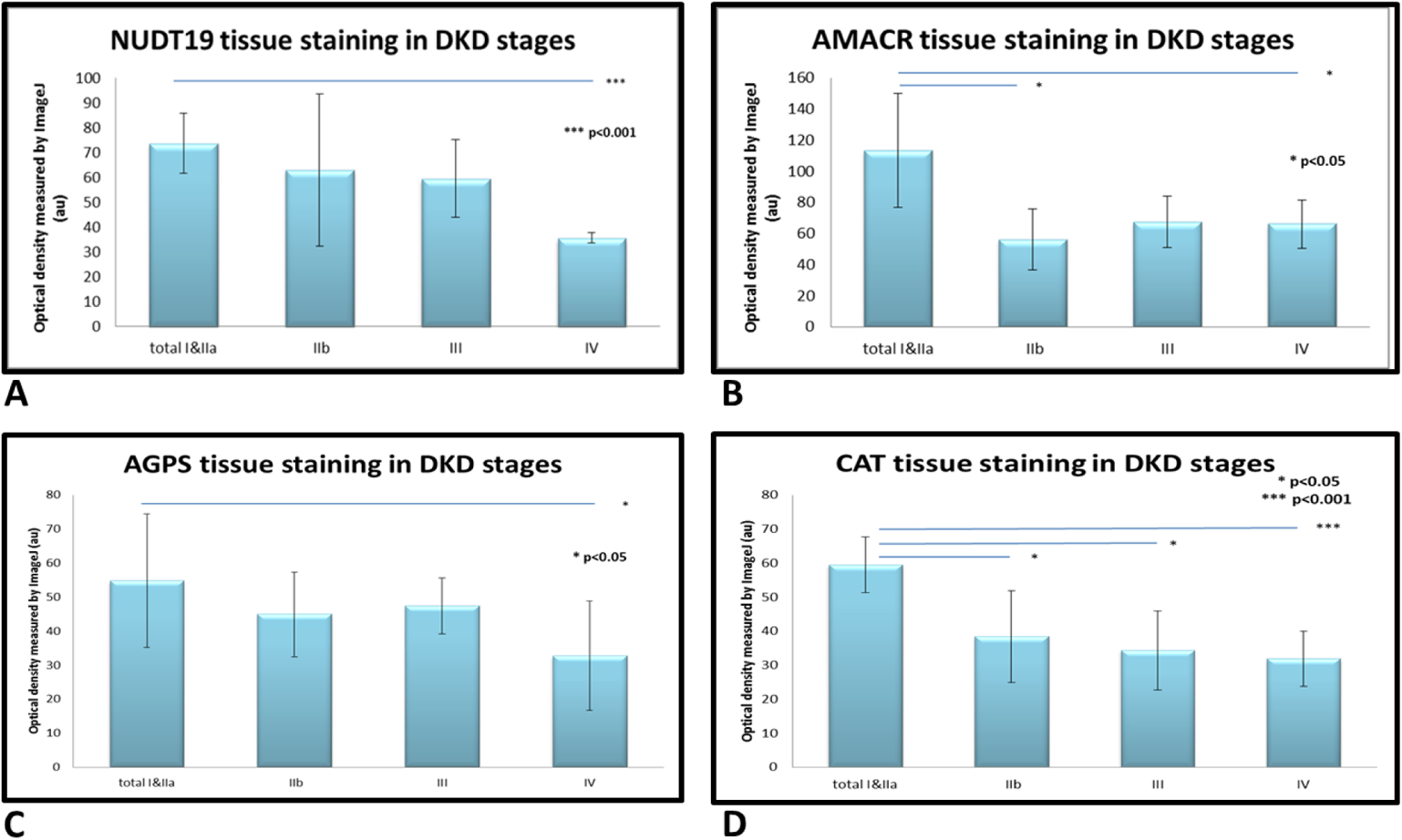
Quantitative analysis of the IHC staining intensity using ImageJ software. Relative quantification scores revealing a decrease in NUDT19 (A), AMACR (B), AGPS (C) and CAT (D) protein expression among groups I+IIa compared to group IV for each of the aforementioned proteins. Values are means ±SD. (n=4 cases of stage I and IIa, 4 cases of stage IIb, 4 cases of stage III and 4 cases of stage IV)(Student’s t-test).

## DISCUSSION

This study was set out with the scope to investigate proteomic changes associated with early DKD and its progression. Proteomic data were generated from two well characterized mouse models (Ins2Akita) of diabetes T1D and (db/db) of T2D. Our study was mainly focused on glomerular proteins that are consistently changed in T1D animals in early and late DKD. The expression of these glomerular proteins was further investigated in the kidney cortex proteome of db/db mice. Multiple consistent changes were observed in proteins involved in cholesterol biosynthesis, mitochondrial respiratory chain function, peroxisomal function and aminoacid metabolism.

The observed elevated levels of mitochondrial enzymes are in agreement with the existing literature: GLS (Glutaminase kidney isoform) catalyzes the first reaction in the kidney catabolism of glutamine (31). Associations of incident prediabetes or T2D with higher levels of glutamate were reported previously (32–36). GLDC (Glycine dehydrogenase-decarboxylating) and AMT (Aminomethyltransferase) participate in mitochondrial glycine cleavage in the kidney (37). Low levels of glycine are related to diabetes and could potentially predict future T2D (36,38–40). Other possible links between low blood-glycine concentration and a metabolic risk are increased glycine utilization for the formation of glutathione to counteract oxidative stress (41) and an increased uptake rate of glycine for gluconeogenesis in insulin-resistant tissues (42). GCAT (2- amino-3-ketobutyrate coenzyme A ligase) participates in the degradation of L-threonine to glycine. Interestingly, decreased threonine levels are reported in diabetes (43). AASS (Alpha-aminoadipic semialdehyde synthase) catalyzes the first two steps in lysine degradation. Of note, lysine levels in the plasma and serum of T2D patients are lower in comparison to controls (33).

The observed changes in peroxisomal proteins, the most prominent observed alteration in our study, are in general agreement with earlier reports (44,45) which suggested decreased expression of key peroxisomal enzymes and regulators of fatty acid oxidation (FAO) in CKD or DKD compared to healthy kidneys.

Peroxisomal enzymes shorten the long chain of very long chain fatty acids (VLCFA), which are subsequently transported to mitochondria and oxidized to acetyl-CoA (46). The first step in peroxisomal FAO is catalyzed by acyl-CoA oxidase (ACOX) and the final product of peroxisomal oxidation can be converted into acyl-carnitine by the carnitine octanoyltransferase (CROT) (47). Interestingly, measurement of CAT, ACOX and CROT in the kidney of ob/ob mice revealed significantly reduced levels (to approximately 2/3) of these peroxisomal enzymes in comparison to controls (48) in line with our results, which is potentially related to the well-established accumulation of lipids in the kidneys of diabetic humans and experimental animals (49,50).

EHHADH (Enoyl-CoA Hydratase And 3-Hydroxyacyl CoA Dehydrogenase) (51) was recently shown to oxidize medium- and long chain fatty acids (52,53). In line with our findings, decreased mRNA level of EHHADH were detected in human DKD glomeruli tissues (54).

Our study also indicated decreased levels of AGPS in mouse proteomics and in human IHC analyses of DKD tissues, with AGPS levels decreasing with DKD progression. AGPS is the main peroxisomal enzyme mediating (bio)synthesis of plasmalogens (55) which act as antioxidants (56), and bile acids. Low levels of plasmalogens have been earlier observed in both T1D (57,58) and T2D (59).

Decreased levels of PIPOX (Peroxisomal sarcosine oxidase) were observed in our diabetic mice. PIPOX lowers pipecolate accumulation through oxidation and increases synthesis of glutaryl-CoA (60,61). In line with our findings, previous studies detected increased levels of pipecolate in T1D mice compared to healthy controls (49). Further, decreased mRNA level of PIPOX were also reported in human DKD glomerular tissues (54).

Our study indicated decreased levels of AMACR, involved in the bile acid biosynthesis pathway (62) in DKD mouse and human kidney tissues. This finding may be linked to earlier observed impairment of bile acid synthesis in T2D (63).

Decreased levels of NUDT19 were also observed in our diabetic mice as well as in human DKD tissues. NUDT19 degrades and regulates CoA in the kidneys (64) thus regulating the peroxisomal CoA pool and b-oxidation (65).

Interestingly, in the cortex of db/db mice of late DKD, 3 additional peroxisomal proteins were detected downregulated: CAT, EPHX2 (Bifunctional epoxide hydrolase 2) and DAO (D-amino acid oxidase; also known as DAAO). CAT, is an antioxidant enzyme (66), whose expression levels also decreased with DKD progression in our human IHC analyses. CAT has also been found downregulated in kidneys and serum from STZ (T2D induced) rodents (67). Interestingly, aberrant catalase activity has been found to increase mitochondrial oxidative stress in kidney proximal tubules (68).EPHX2 (an antioxidant enzyme) protein and mRNA levels, in accordance with our study, were found decreased in the kidneys of streptozotocin (STZ)-induced diabetic mice (69) as well as rodents of progressive kidney disease (70). Finally, DAO-participating in amino acid degradation (71) - expression was also decreased in the kidney of DKD alloxan-diabetic rats, in line with our results (72).

Collectively, our study highlights and brings together multiple protein changes, which had to a good extent been observed at the mRNA level in animal models or human tissue, supporting, as a step further, their role at an early time point in DKD development. Based on our results, a massive disruption of the mitochondrial-peroxisome cross talk in early DKD is supported (summarized in Figure 7) with special emphasis on fatty acid oxidation (in which most of the reported dysregulated proteins participate). This is of great interest since most of the peroxisome studies in DKD are focused mainly on the mitochondrial oxidative stress and the role of peroxisomal catalase (66). Although there are no drugs so far targeting any of the detected peroxisomal proteins, it is observed that SGLT2i reduce oxidative stress in the kidney of mouse diabetic models (db/db, Ins2Akita and ob/ob) (80–82) As a next step we could validate our results in a large cohort of human DKD kidney samples in order to unveil the molecular details of mitochondrial-peroxisomal cross talk in DKD which remain unclear.

**Figure 7.**
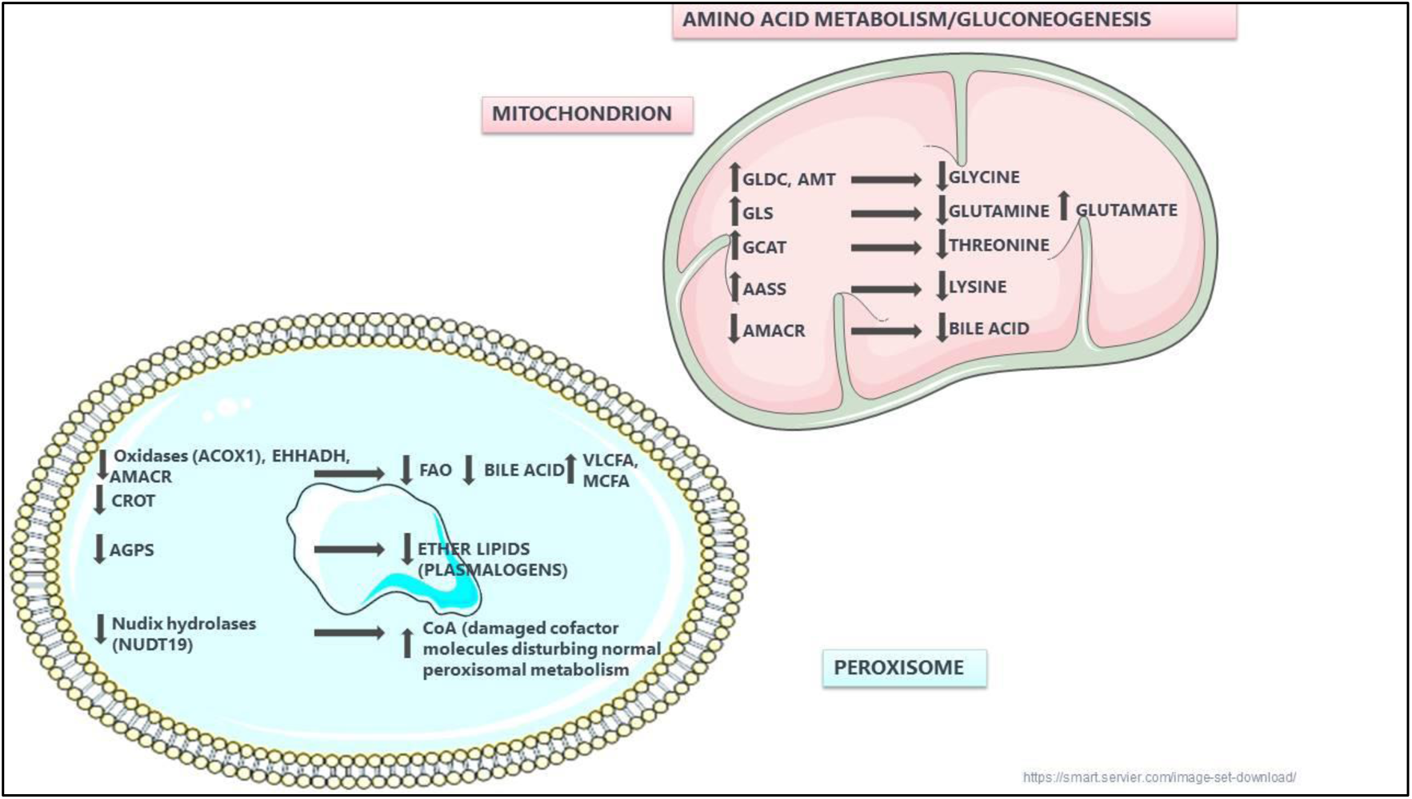
Peroxisomal and mitochondrial dysfunction in DKD based on consistently changed proteins in our study. It may be hypothesized that the observed decreased expression of peroxisomal proteins involved in lipid metabolism and CROT leads to elevated levels of VLCFA and MCFA, reduced bile acid synthesis and disruption of acylcarnitines transportation to mitochondria. Moreover, downregulation of AGPS reduces plasmalogens synthesis. Peroxisomal Nudix hydrolases, such as NUDT19 (hydrolyzes CoA), clean the cell from harmful metabolites such as ROS indicating their role as ‘housecleaning’ enzymes. In mitochondria increased expression of GLDC, GLS, GCAT and AASS favors gluconeogenesis.

## Acknowledgments

This project has received funding from the Hellenic Foundation for Research and Innovation (HFRI) and the General Secretariat for Research and Innovation (GSRI), under grant agreement No 695 (MolProt-CKD).

## Ethics approval and consent to participate

All animal experiments were conducted in accordance with the Guide for the care and use of laboratory animals of the National Institute of Health, eighth edition and the French Institute of Health guidelines for the care and use of laboratory animals. The project was approved by the local (Inserm/UPS US006 CREFRE) and national ethics committees (ethics committee for animal experiment, CEEA122; Toulouse, France; approval 02867.01 for Ins2Akita mice and 690966 for db/db mice).

The study of human samples was approved by the National and Kapodistrian University of Athens Medical School Ethics Committee; since this was a retrospective study, the Ethics Committee waived the need for an informed consent, and a policy of strict anonymity and confidentiality was assured.

## Data sharing

Anonymised data will be made available upon request directed to the corresponding author. Proposals will be reviewed and approved by the investigators and collaborators based on scientific merit. After approval of a proposal, data will be shared through a secure online platform after signing a data access and confidentiality agreement.

## Supplementary Material

**Figure S1**

Kinetics of weight, glycemia and ACR in Ins2Akita and db/db mice. Blood and urine of controls (wt and db/dm respectively) and Ins2Akita (A) and db/db (B) mice were collected at different age, and weight, glycemia and ACR were quantified. Values are mean ± SEM.

**Figure S2**

Correlation analysis of the protein abundance between our dataset and the study of Waanders LF et al 2009 (22) (PMID 19846766). A correlation factor of R=0.7 for the wild type mice and R=0.7 for the Ins2Akita mice was observed.

**TableS1**

Summary of DKD datasets extracted from Nephroseq database.

**TableS2**

The main clinicopathological parameters of the 16 human cases of IHC analysis.

**TableS3**

The full list of all proteomics data of each sample of wild type and Ins2Akita mice 2 and 4 month old respectively.

**TableS4**

The full list of identified proteins using a 55% frequency threshold of wild type and Ins2Akita mice 2 and 4 month old respectively including the fold change (ratio) and the statistics.

**TableS5**

The biological process of the upregulated and downregulated proteins comparing Ins2akita vs WT of 2 month and 4 month old respectively.

**TableS6**

Statistically significant changed proteins comparing WT4 month old vs WT2 month old.

**TableS7**

List of the glomerular proteins consistently changed in early and late DKD versus controls which are related to diabetes and/or have attributed kidney expression and/or function.

**TableS8**

The full list of all proteomics data of each sample of wild type and db/db mice 6 month old respectively.

**TableS9**

A comprehensive list of the identified proteins using a 60% frequency threshold of wild type and db/db mice 6 month old respectively including the fold change (ratio) and the statistics.

**TableS10**

Cross-omics validation of deregulated consistently changed mitochondrial and peroxisomal glomerular proteins in early and late DKD. The upregulated proteins are presented in green and the downregulated in red color.

**TableS11**

Cross-omics validation of additional peroxisomal deregulated proteins detected in the cortex collected from db/db mice of late DKD. The downregulated proteins are presented in red color.

