## Supplementary figures and images for "Proteomic analysis of mouse kidney tissue associates peroxisomal dysfunction with early diabetic kidney disease"

### FigureS1

**Figure S1**

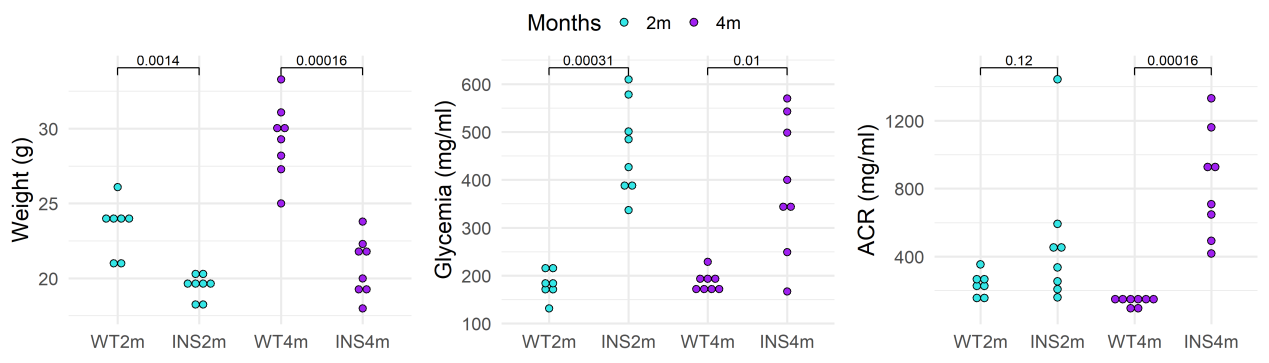

A)

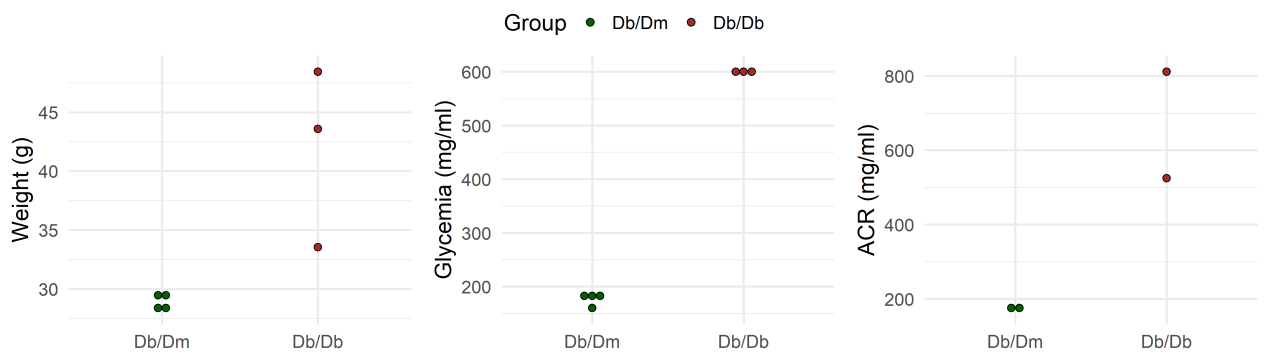

Β)

### FigureS2

**Figure S2**

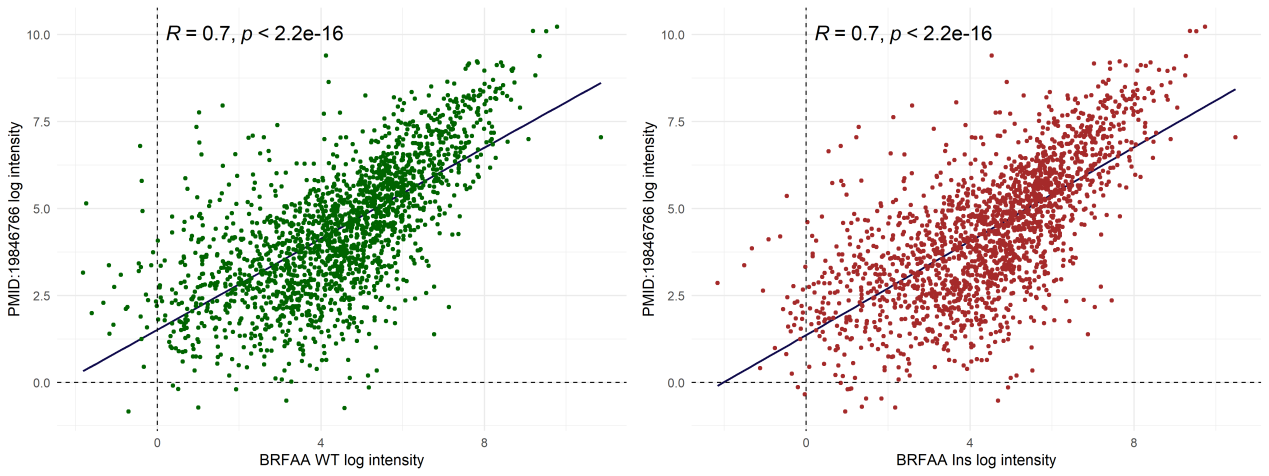
