## Supplementary material for "Proteomic analysis of mouse kidney tissue associates peroxisomal dysfunction with early diabetic kidney disease": TableS2

| **Case No** | **Gender** | **Age** | **DM**  **history** | **DKD**  **Class** | **IFTA score** | **Inflammation score** | **Hyalinosis score** | **Art/sclerosis score** | **Total score** | **CKD**  **Stage** | **SCr**  **(mg/dL)** | **Albuminuria (g/24h)** | **eGFR**  **ml/min/1,73 m^2^** |
| --- | --- | --- | --- | --- | --- | --- | --- | --- | --- | --- | --- | --- | --- |
| 1 | M | 51 | DM type II | **I** | 1 | 0 | 2 | 1 | 4 | **G1A3** | 0.9 | 0.5 | 99 |
| 2 | M | 55 | DM type II | **IIa** | 2 | 1 | 2 | 1 | 6 | **G1A3** | 0.8 | 4 – 6 | 101 |
| 3 | M | 53 | DM type II | **IIa** | 2 | 1 | 2 | 1 | 6 | **G1A3** | 0.95 | 5.4 | 91 |
| 4 | M | 50 | DM type II | **IIa** | 2 | 1 | 2 | 1 | 6 | **G2A3** | 1.2 | 1.8 | 70 |
| 5 | M | 82 | DM type II | **IIb** | 3 | 2 | 2 | 2 | 9 | **G2A3** | 1 | ≥3 | 70 |
| 6 | M | 30 | DM type I | **IIb** | 3 | 1 | 2 | 1 | 7 | **G3aA3** | 1.7 | 8 | 53 |
| 7 | F | 35 | DM type I | **IIb** | 3 | 1 | 2 | 2 | 8 | **G3aA3** | 1.3 | 16.8 | 53 |
| 8 | M | 73 | DM type II | **IIb** | 2 | 1 | 2 | 2 | 7 | **G3bA3** | 1.6 | ≥3 | 42 |
| 9 | M | 58 | DM type II | **III** | 3 | 1 | 2 | 2 | 8 | **G3bA3** | 2 | 3 | 36 |
| 10 | M | 80 | DM type II | **III** | 3 | 1 | 2 | 2 | 8 | **G3bA3** | 1.9 | ≥3 | 33 |
| 11 | M | 69 | DM type II | **III** | 2 | 1 | 2 | 0 | 5 | **G3bA3** | 2 | 4 | 33 |
| 12 | M | 58 | DM type II | **III** | 3 | 1 | 2 | 2 | 8 | **G3bA3** | 2.2 | ≥3 | 32 |
| 13 | M | 65 | DM type II | **IV** | 3 | 1 | 2 | 2 | 8 | **G4A3** | 2.9 | 2 | 22 |
| 14 | F | 62 | DM type I | **IV** | 3 | 1 | 2 | 2 | 8 | **G4A3** | 2.5 | 0.6 | 20 |
| 15 | M | 50 | DM type II | **IV** | 3 | 1 | 2 | 2 | 8 | **G5A3** | 5.5 | ≥3 | 11 |
| 16 | M | 44 | DM type I | **IV** | 3 | 1 | 2 | 2 | 8 | **G5A3** | 7 | 10 | 9 |
