## Supplementary material for "Proteomic analysis of mouse kidney tissue associates peroxisomal dysfunction with early diabetic kidney disease": TableS10, 11

| **Protein** | **Transcriptomics expression (Nephroseq; in DKD vs. controls) (ref)** | **Single-cell human kidney transcriptomics expression (Wilson et al. 29)** | **Protein expression or metabolites (ref)** |
| --- | --- | --- | --- |
| **GLS** | increase (26,27) | increase (tubuli) | No proteomics data only on metabolites (31-36) reduced levels of glutamine and/or elevated levels of glutamate in diabetes |
| **GLDC** | increase (28 db/db mouse glomeruli) | increase (tubuli) | No proteomics data only on metabolites (36-40) reduced levels of glycine in diabetes |
| **AMT** | decrease (26) | not detected | No proteomics data only on metabolites (36-40) reduced levels of glycine in diabetes |
| **ACOX1** | no data | decrease (tubuli) | decrease (48,49) |
| **CROT** | decrease (28 db/db mouse glomeruli) | decrease (glomeruli) | decrease (48) ob/ob |
| **EHHADH** | decrease (27, 28 db/db mouse glomeruli) | decrease (glomeruli) | not detected |
| **AGPS** | decrease (28 db/db mouse glomeruli) | decrease (glomeruli) | No proteomics data only on metabolites (56-58) low plasmalogens in diabetes |
| **PIPOX** | decrease (27) | not detected | No proteomics data only on metabolites (45, 63) elevated pipecolate in diabetes |
| **AMACR** | decrease (27) | decrease (glomeruli) | No proteomics data only on metabolites (63) reduced levels of bile acid synthesis in diabetes |
| **NUDT19** | decrease (28 db/db mouse glomeruli) | not detected | not detected |
| **PECR** | decrease (28 db/db mouse glomeruli) | not detected | not detected |

**TableS11**

| **Protein** | **Transcriptomics expression (Nephroseq; in DKD vs. controls) (ref)** | **Single-cell human kidney transcriptomics expression (Wilson et al. 29)** | **Protein expression (ref)** |
| --- | --- | --- | --- |
| **CAT** | decrease (28 db/db mouse glomeruli) | decrease (tubuli) | decrease (67) STZ mice |
| **EPHX2** | no data | decrease (tubuli) | not detected |
| **DAO** | decrease (26,27) | decrease (glomeruli and tubuli) | decrease (72) |
